# The Arabidopsis BUB1/MAD3 family protein BMF3 requires BUB3.3 to recruit CDC20 to kinetochores in spindle assembly checkpoint signaling

**DOI:** 10.1101/2023.06.22.545541

**Authors:** Xingguang Deng, Felicia Peng, Xiaoya Tang, Yuh-Ru Julie Lee, Hong-Hui Lin, Bo Liu

## Abstract

Mitosis is monitored by the spindle assembly checkpoint (SAC) which remains active until all chromosomes have their kinetochores attached to microtubules originated from opposite spindle poles. Plants produce both highly conserved and sequence-diverged SAC components, so it has been largely unknown how SAC activation leads to the assembly of these proteins at the unattached kinetochores to prevent anaphase onset in plants. In *Arabidopsis thaliana*, the noncanonical BUB3.3 protein was detected at kinetochores throughout mitosis, unlike MAD1 and the plant specific BUB1/MAD3 family protein BMF3 that associated with unattached chromosomes only. However, BUB3.3 was required to arrest mitotic progression when one or more chromosomes did not congress at the metaphase plate. Nevertheless, BUB3.3 was not required for the kinetochore localization of other SAC components and *vice versa*. BUB3.3 specifically interacted with BMF3 in a novel region containing two internal repeats that were not required for the kinetochore localization of BMF3. This interaction was important for BMF3 to recruit the CDC20 protein to the unattached kinetochores. Our results showed that the Arabidopsis BUB3.3 protein functioned in the activation of BMF3 for CDC20 recruitment, rather than the recruitment of BMF proteins as what has been found in fungi and animals, in order to arrest mitosis at prometaphase. Therefore, activated SAC resulted in BUB3.3-independent localization of BMF3 and MAD1 to kinetochores and BUB3.3-dependent licensing of BMF3 for CDC20 recruitment in *A*. *thaliana*.

**SIGNIFICANCE STATEMENT:** Mitotic progression into anaphase is monitored by spindle assembly checkpoint that is poorly understood in plants. Using *Arabidopsis thaliana* as a reference system, we discovered a novel interaction pattern centered at the BUB1 and MAD3 protein BMF3 at kinetochores. A noncanonical isoform of the evolutionarily conserved BUB3 family protein BUB3.3 interacted with two novel internal repeats in BMF3 for recruiting the CDC20 protein to unattached kinetochores in order to inhibit its function in anaphase onset. Hence, our work sheds light on how spindle assembly checkpoint operates in flowering plants that produce the highly conserved BUB3.3 protein but highly divergent BMF proteins.

## INTRODUCTION

Cell division results in the production of two daughter cells with equal amounts of genetic materials. Faithful karyokinesis requires all chromosomes to be aligned at the metaphase plate, resulted from the attachment of sister kinetochores to microtubule fibers originated from opposite poles, prior to anaphase onset. Monitoring such an amphitelic chromosome attachment or biorientation is the spindle assembly checkpoint (SAC) that is dependent on what are called BUB (Budding Uninhibited by Benzimidazoles) and MAD (Mitotic Arrest Defective) proteins (1). In fungi and animals, SAC activation results in the kinase-dependent recruitment of MAD1 and MAD2 to kinetochores where MAD2 of the closed conformation is converted from the prior open one. Such closed MAD2 is joined by BUBR1/MAD3 which is docked on the kinetochores by the WD40 repeat protein BUB3 to form the mitotic checkpoint complex (MCC) consisting of BUB3, BUBR1/MAD3, MAD2, and CDC20 which is the activator of the APC/C (Anaphase Promoting Complex/Cyclosome). MCC-bound APC/C prevents anaphase onset from taking place so that mitosis is arrested due to SAC activation (1).

Flowering plants like *Arabidopsis thaliana* produce BUB3, MAD1, and MAD2 homologs resembling their counterparts in fungi and animals, as well as three proteins distantly related to BUB1 and BUBR1/MAD3 (2). These three so-called BUB1/MAD3 family proteins or BMFs all possess an BUB1/MAD3-type TPR (tetratrichopeptide repeat) domain but otherwise share little if any homology outside this N-terminal domain (2). Only BMF1, but not BMF2 and BMF3, has a C-terminal kinase domain, and none of the BMF proteins possess the BUB3-interacting GLEBS (Gle2-binding sequence) domain found in fungal and animal BUB1 and BUBR1/MAD3 proteins (3, 4). Therefore, it is unknown whether the BUB3 protein interacts with BMFs or whether it is a *bona fide* SAC component that acts at the kinetochores when the SAC is turned on.

The hallmark feature of SAC components is their localization at the kinetochores of unattached chromosomes and dissociation from metaphase plate-aligned chromosomes, as often demonstrated in animal cells (1). In *A*. *thaliana*, MAD1 and BMF3 exhibit such featured kinetochore localization prior to reaching metaphase but BMF1 resides at kinetochores throughout mitosis (4). In contrast, other SAC proteins exhibit different localization patterns during mitosis, for example, BMF2 was undetectable by using a functional BMF2-GFP (green fluorescent protein) fusion and a functional MAD2-GFP fusion was most abundant in a diffuse pattern in the cytoplasm (4). There are three BUB3 proteins among which BUB3.1 and BUB3.2 are closely related to the fungal and animal BUB3 but are associated with phragmoplast microtubules by directly interacting with the microtubule-associated protein MAP65-3 for cytokinesis (5). In contrast, it is unclear where the less conserved, noncanonical BUB3.3 protein may function during mitosis.

The characteristic phenotype of the loss of an essential SAC component is the hypersensitivity to microtubule-depolymerizing drugs like the herbicide oryzalin commonly used in plant studies. For example, the *bub3.3*, *mad1*, *mad2*, and *bmf3* mutants grow indistinguishable from the wild-type control but had their root growth almost completely inhibited by 150 nM oryzalin that does not obviously harm the growth of wild-type plants (4). The *bmf1* mutant is insensitive to oryzalin and the *bmf2* mutant is less sensitive than *bmf3*, suggesting that BMF1-dependent phosphorylation is not required for SAC regulation and BMF2 either plays a weak role or is not involved in SAC (2). Thus, the differences in the protein architecture, localization, and function of the plant MAD and BMF proteins from those in fungi and animals raised the question of how plants engage necessary SAC components in the assembly of the MCC-like complex when the SAC is turned on.

The oryzalin hypersensitivity phenotype of *bub3.3* prompted us to test whether the MAD1 and BMF3 function at kinetochores upon SAC activation was dependent on BUB3.3, like what has been described in fungi and animals. Fungal and animal BUB3 proteins recognize GLEBS or MELT motif found in their interacting proteins (6). But none of the BMF proteins contains these motifs, thus raised the question of whether BUB3.3 directly interacted with one or more of them and whether it acted in concert with BMF3. To gain insights into SAC regulation in *A*. *thaliana*, we investigated the function of BUB3.3 and discovered a novel mode of interaction between BUB3.3 and BMF3 for the recruitment of CDC20 to unattached kinetochores in order to prevent anaphase onset from taking place before all chromosomes were aligned at the metaphase plate in mitotic cells.

## RESULTS

### BUB3.3 associates with kinetochores throughout mitotic cell division

To examine how BUB3.3 functioned during mitosis, we transformed a *bub3.3* mutant with a construct for GFP-BUB3.3 expression under the control of the BUB3.3 promoter. When we tested the transformants together with the wild-type control and the *bub3.3* mutant, we found that the *bub3.3* mutant showed significant growth retardation when exposed to 100 nM oryzalin, as quantified by the root lengths (Figure 1A, B). Such an oryzalin hypersensitivity phenotype was suppressed when the *BUB3.3*(*p*)::GFP-*BUB3.3* transgene was expressed in the homozygous mutant (Figure 1A, B), indicating that the phenotype was caused by the inactivation of the *BUB3.3* gene.

**Figure 1.**
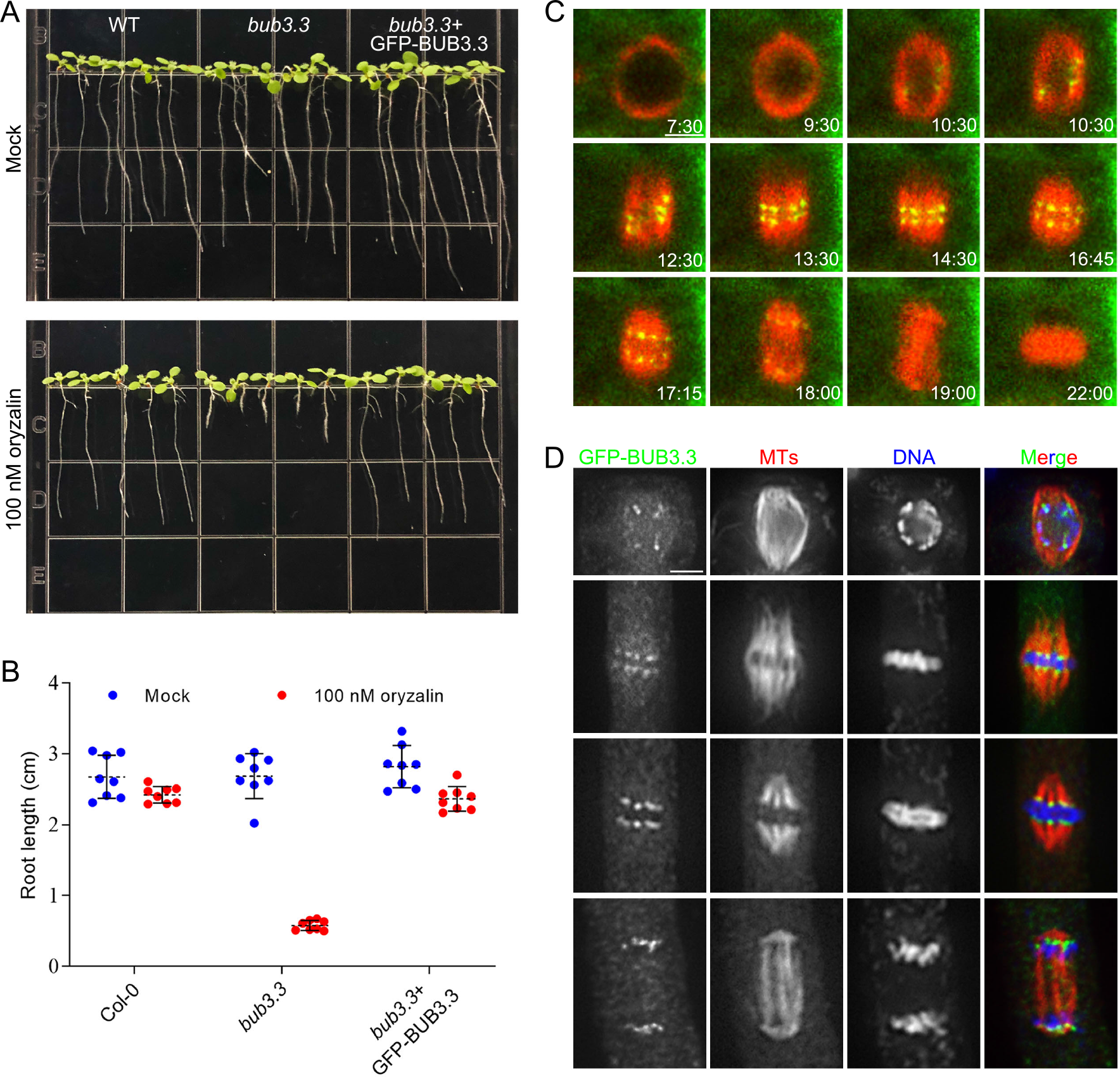
BUB3.3 associates with kinetochores throughout mitotic cell division. (**A**) Ten-day-old seedlings grown in the absence of presence of 100 nM oryzalin. The wild-type (WT) control, *bub3.3* mutant, and *bub3.3* mutant expressing GFP-BUB3.3 are included. (**B**) Quantification of root lengths of the seedlings with and without oryzalin treatment (n= 8 for each sample). (**C**) Live imaging of GFP-BUB3.3 and mCherry-TUB6 (for microtubules) in a mitotic cell. Snapshots are taken from Supplementary Movie S1. (**D**) Triple localization of BUB3.3, microtubules, and DNA mitosis. The merged images have GFP-BUB3.3 detected by the anti-GFP antibody pseudocolored in green, microtubules in red and DNA in blue. Scale bars = 5 μm.

Because the GFP-BUB3.3 fusion protein was functional as indicated by genetic suppression of the *bub3.3* mutation, we used the transgenic line to examine subcellular localization of GFP-BUB3.3 during mitosis. An mCherry-TUB6 (β-tubulin 6) fusion was introduced into the transgenic plant for us to monitor mitotic progression in live-cells. GFP-BUB3.3 was detected as paired dots inside the prophase nucleus surrounded by a bipolar microtubule array (Figure 1C). The signal became aligned in the middle of the spindle before being separated into two groups towards opposite spindle poles in the cell. The prominent signal remained in bright dots at telophase (Figure 1C). To determine the relationship between the GFP-BUB3.3 signal and kinetochores/chromosomes, we performed immunofluorescence experiments in fixed cells. At late prophase when a bipolar spindle array was formed, separated GFP-BUB3.3 dots were detected in the nucleus that were associated with chromosomes (Figure 1D). At metaphase when chromosomes were aligned at the metaphase plate that was flanked by mirrored kinetochore fibers, the GFP-BUB3.3 signal appeared at the two edges of aligned chromosomes and the end of kinetochore fibers (Figure 1D). When kinetochore fibers shortened, the GFP-BUB3.3 signal followed the shortening fibers until they reached the two spindle poles at telophase (Figure 1D). Therefore, this dynamic BUB3.3 localization led to the conclusion that it was associated with kinetochores throughout mitosis.

### The loss of BUB3.3 causes frequent chromosomes misalignment

BUB3 is central for BUB1 and BUBR1 to exercise their functions at the kinetochores in fungi and animals (1). Because BUB3.3 is the kinetochore localized BUB3 protein in *A*. *thaliana*, unlike the phragmoplast-associated BUB3.1 and BUB3.2, we then asked whether its loss led to defects in mitosis by examining meristematic cells. Although the *bub3.3* mutant produced healthy plants, its mitotic cells often exhibited phenotypes of chromosome misalignment (Figure 2A). In wild-type cells producing typical metaphase spindles with paired kinetochore microtubule fibers without oryzalin treatment, chromosomes were always aligned at the metaphase plate (Figure 2A). In the *bub3.3* cells, however, mitotic cells bearing similar spindle microtubule arrays often had one or two chromosomes left outside the metaphase plate, towards one side of the cells, and other cells had a misaligned chromosome on each side of the cells, near two spindle poles (Figure 2A). In fact, over 60% of the mitotic cells had misaligned chromosomes outside the metaphase plate where the majority of chromosomes had been congressed (Figure 2D). After the cells were treated with 100 nM oryzalin, greater than 95% (62/65) of the *bub3.3* cells had exacerbated chromosome misalignment and misaligned chromosomes sometimes outnumbered ones at the metaphase plate (Figure 2B, D). Furthermore, the misaligned chromosomes typically had microtubules attached but separated from the main spindle that had congressed chromosomes engaged as if a “minispindle” was formed outside the main one (Figure 2C). Such a challenge also caused subtle chromosome misalignment phenomena in the control cells (Figure 2B, D). We then examined whether the misaligned chromosomes were sufficient to give birth to a nucleus structure by examining cytokinetic cells. In wild-type cells, cytokinesis always resulted in the birth of two identically looking nuclei with or without oryzalin treatment (Figure 2E, F). In the *bub3.3* mutant, micronuclei were formed towards the end of cytokinesis after oryzalin treatment that were otherwise not observed without oryzalin (Figure 2E, F). Therefore, the results suggested that the misaligned chromosomes in the mutant cells were able to join others to form daughter nuclei and exacerbation of the misalignment by oryzalin treatment enhanced the phenotype so that these left-out chromosomes reformed micronuclei.

**Figure 2.**
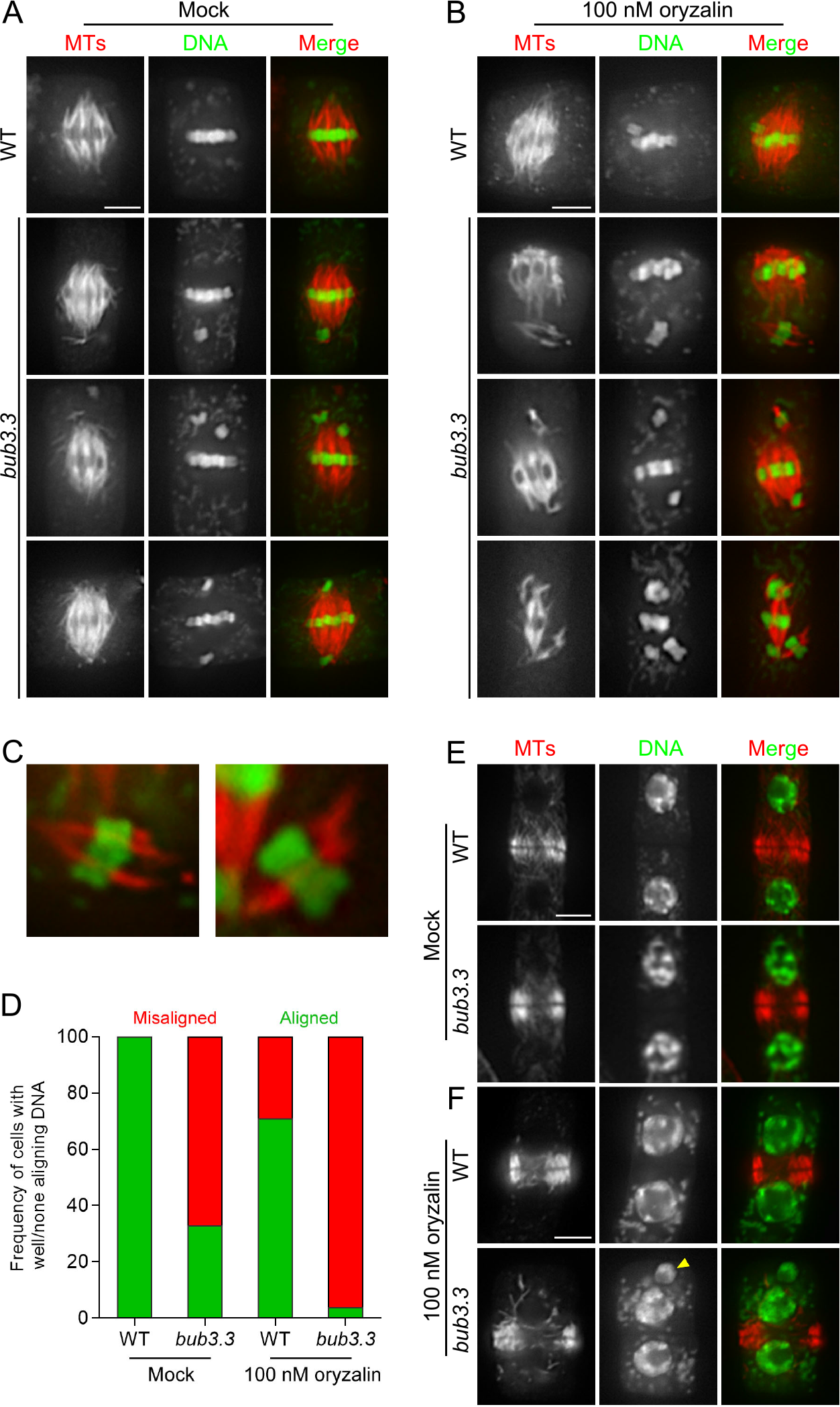
Chromosomes misalignment in the absence of BUB3.3. (**A, B**) Chromosome alignment in wild-type (WT) control and *bub3.3* mutant cells at late stages of prometaphase in the absence (**A**) or presence (**B**) of oryzalin. (**C)** Enlarged views of kinetochore fibers/minispindles with misaligned chromosomes in *bub3.3* cells. (**D)** Quantitative assessment of abnormal cells exhibiting misaligned chromosomes in *bub3.3* cells compared to WT cells with or without oryzalin treatment (n=65). (**E, F)** Comparative views of cytokinetic cells in WT and *bub3.3* plants in the absence (**E**) or presence (**F**) of oryzalin. The yellow arrowhead points at a representative micronucleus formed after oryzalin treatment in *bub3.3* cells. Merged images have microtubules in red and DNA in green. Scale bars = 5 μm.

### The *bub3.3* mutant cells can enter anaphase with misaligned chromosomes in the presence of oryzalin

The oryzalin hypersensitivity phenotype prompted us to test whether the loss of BUB3.3 led to errors in mitosis. To monitor mitotic progression in living cells, we delivered a NDC80-TagRFP marker that labeled kinetochores, like what has been reported (7). When wild-type roots were exposed to 100 nM oryzalin, we detected the NDC80-TagRFP signal outside the metaphase plate, representing a misaligned chromosome (Figure 3A). Anaphase took place after this misaligned chromosome was brought to the metaphase plate (arrowheads, Figure 3A. Supplemental Movie 2). In the *bub3.3* mutant cells after the oryzalin treatment, mitotic cells entered anaphase without having the misaligned chromosome congress to the metaphase plate (arrowheads, Figure 3B. Supplemental Movie 3). Consequently, unaligned chromosome(s) joined the segregated sister chromatids so that erroneous mitosis took place because of unequal chromosome segregation (Figure 3B). Therefore, our result provided firsthand evidence showing that the loss of this noncanonical BUB3.3 led to failures in activating the SAC activation in response to misaligned chromosomes.

**Figure 3.**
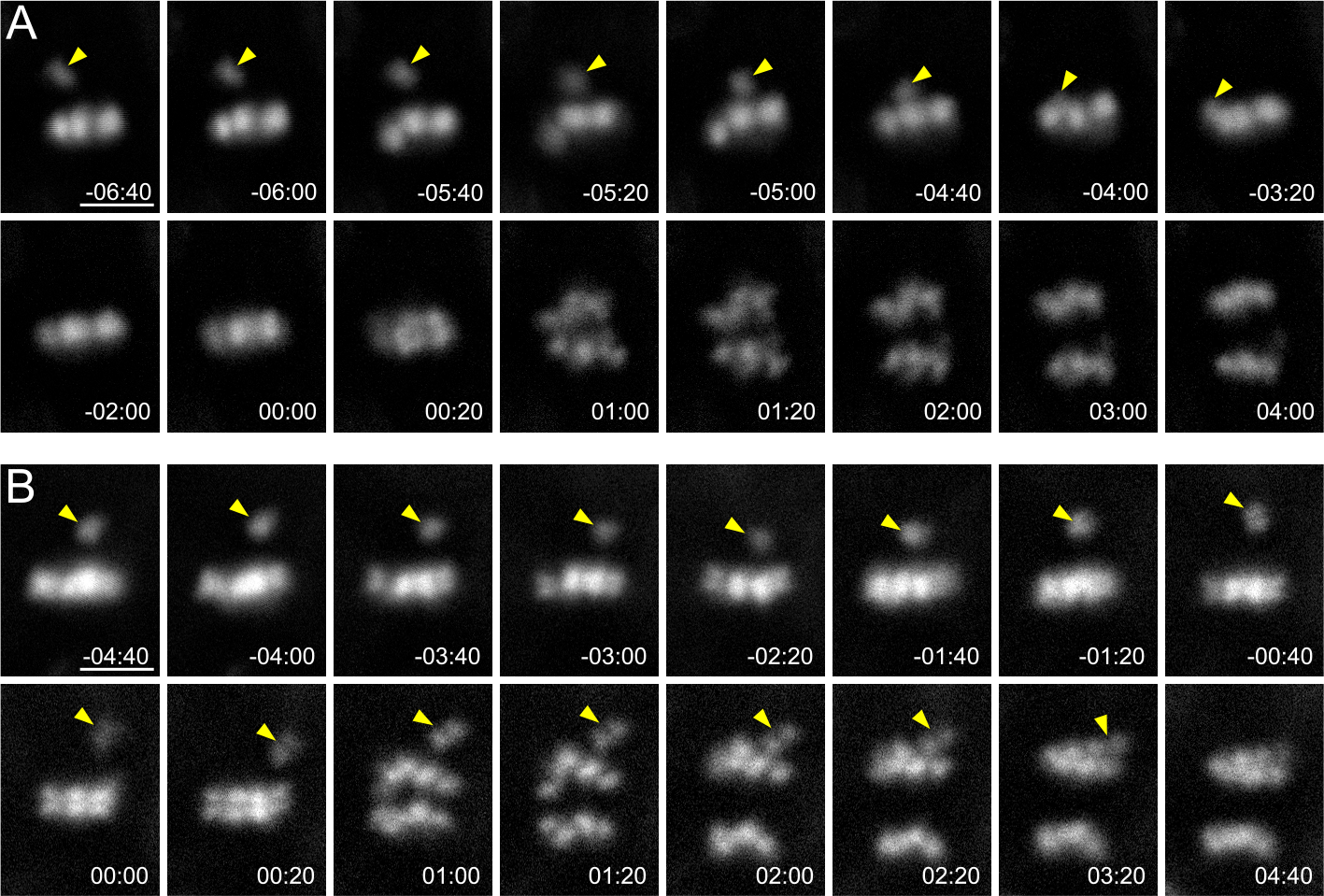
BUB3.3 monitors chromosome alignment at the metaphase plate to prevent premature sister chromatid segregation. (**A, B)** Mitotic progression is monitored by live imaging of the NDC80-TagRFP fusion protein in WT (**A**) and *bub3.3* (**B**) cells treated with 100 nM oryzalin. Snapshots are taken from Supplemental Movies 2 and 3. In the WT cell, anaphase onset takes place after the misaligned chromosome (arrowhead) is brought to the metaphase plate. In *bub3.3* cells, however, the cell ignores the misaligned chromosome (arrowhead) and enters anaphase. Scale bars = 5 μm.

### BUB3.3 localization at kinetochores is independent to other SAC proteins

Because BUB3 is a key targeting factor for the kinetochore localization of MAD and BUB proteins in SAC activated animal cells, we tested whether such a role was shared by the BUB3.3 protein in *A*. *thaliana*. To do so, we transformed constructs, which have been demonstrated to be functional GFP fusion proteins of MAD and BMF proteins (4), into the *bub3.3* mutant. The BMF1-GFP fusion protein decorated kinetochores throughout mitosis in both the control and the *bub3.3* mutant (Figure 4A, B). BMF1-GFP did not distinguish metaphase plate aligned or misaligned (arrowheads) chromosomes in the *bub3.3* mutant (Figure 4B). Unlike BMF1, MAD1 and BMF3 are removed from the kinetochores once chromosomes are aligned at the metaphase plate and the SAC is turned off (Figure 4C, E) (4). Such a feature was preserved in the *bub3.3* mutant in which prominent GFP-MAD1 and BMF3-GFP signals were detected at the kinetochores prior to chromosome alignment or at those of frequently misaligned chromosomes (arrowheads) while aligned ones lacked the signals (Figure 4D, F).

**Figure 4.**
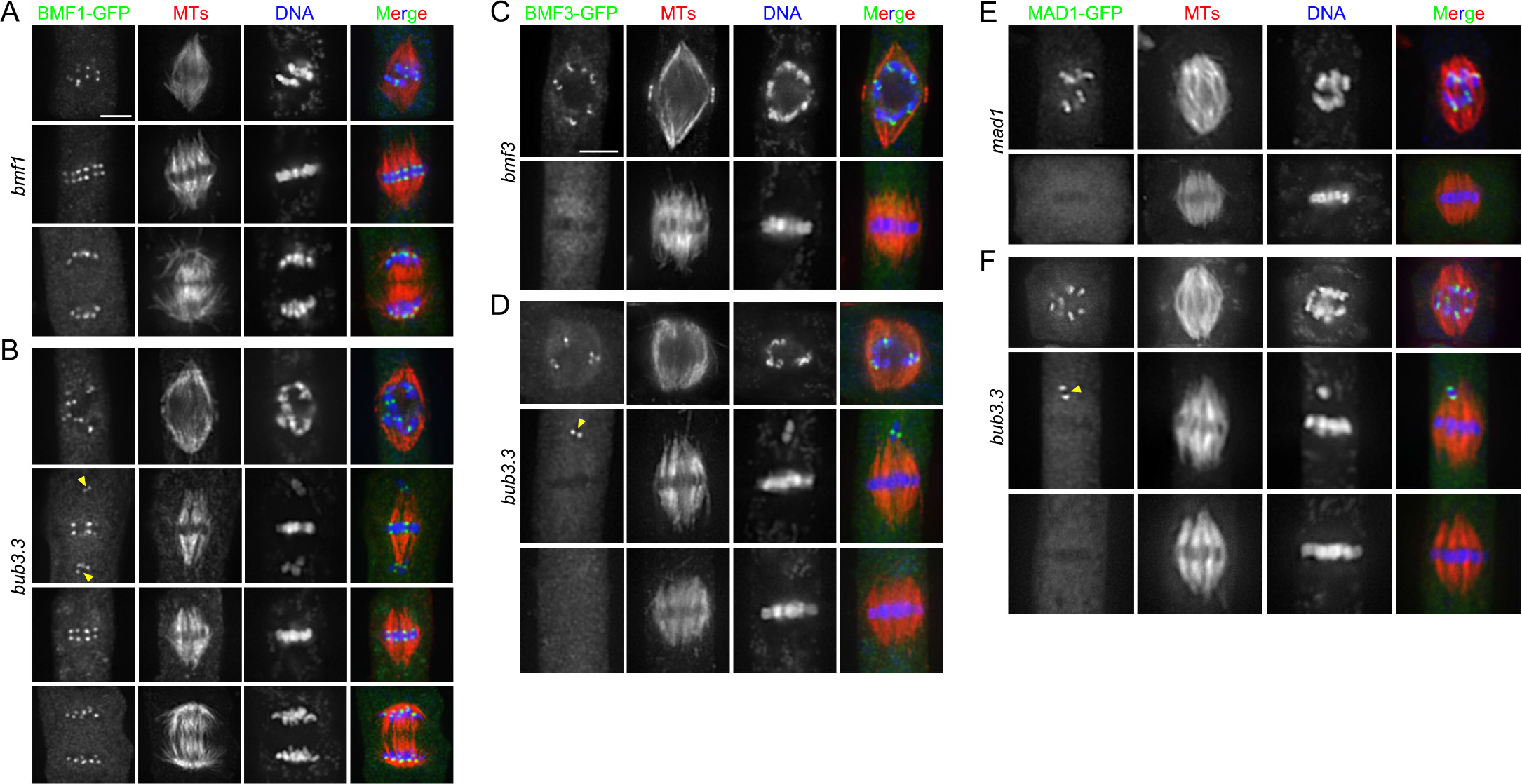
BUB3.3 is not required for the kinetochore localization of other SAC proteins. (**A, B)** The BMF1-GFP fusion protein localizes to kinetochores of both aligned and misaligned chromosomes in *bmf1* (**A**) and *bub3.3* (**B**) mutant cells. (**C, D)** BMF3-GFP decorates kinetochores of unattached/misaligned chromosomes similarly in *bmf3* (**C**) and *bub3.3* (**D**) mutant cells. (**E, F)** MAD1-GFP, like BMF3, only localizes to kinetochores of unaligned chromosomes in both *mad1* (**G**) and *bub3.3* (**H**) mutant cells. Note that in the *bub3.3* mutant cells, the BMF3 and MAD1 signals are no longer detected at the kinetochores when chromosomes arrive at the metaphase plate. Merged images have GFP-tagged proteins in green, microtubules in red, and DNA in blue. Scale bars = 5 μm.

Conversely, we asked whether any of the MAD and BMF proteins were required for BUB3.3 localization. When GFP-BUB3.3 was expressed in the *bmf1* and *mad1* mutant, it was associated with kinetochores at all stages of mitosis (Figure 5A, D), as demonstrated earlier (Figure 1C). Because BMF2 and BMF3 but not BMF1 have been detected as functional SAC components based on the oryzalin hypersensitivity test (4), we generated a *bmf2*; *bmf3* double mutant and expressed the GFP-BUB3.3 fusion in this background. Again, GFP-BUB3.3 was undisturbed at the kinetochores throughout mitosis (Figure 5B). Finally, we checked the *mps1* mutation because it shows the oryzalin hypersensitivity phenotype like other sensitive *bmf* mutants (4). Like other mutations, the loss of MPS1 did not affect GFP-BUB3.3 localization either (Figure 5C). Therefore, our results indicated that BUB3.3 localization is independent to the other critical SAC proteins of MPS1, MAD1, BMF2, and BMF3, and vice versa.

**Figure 5.**
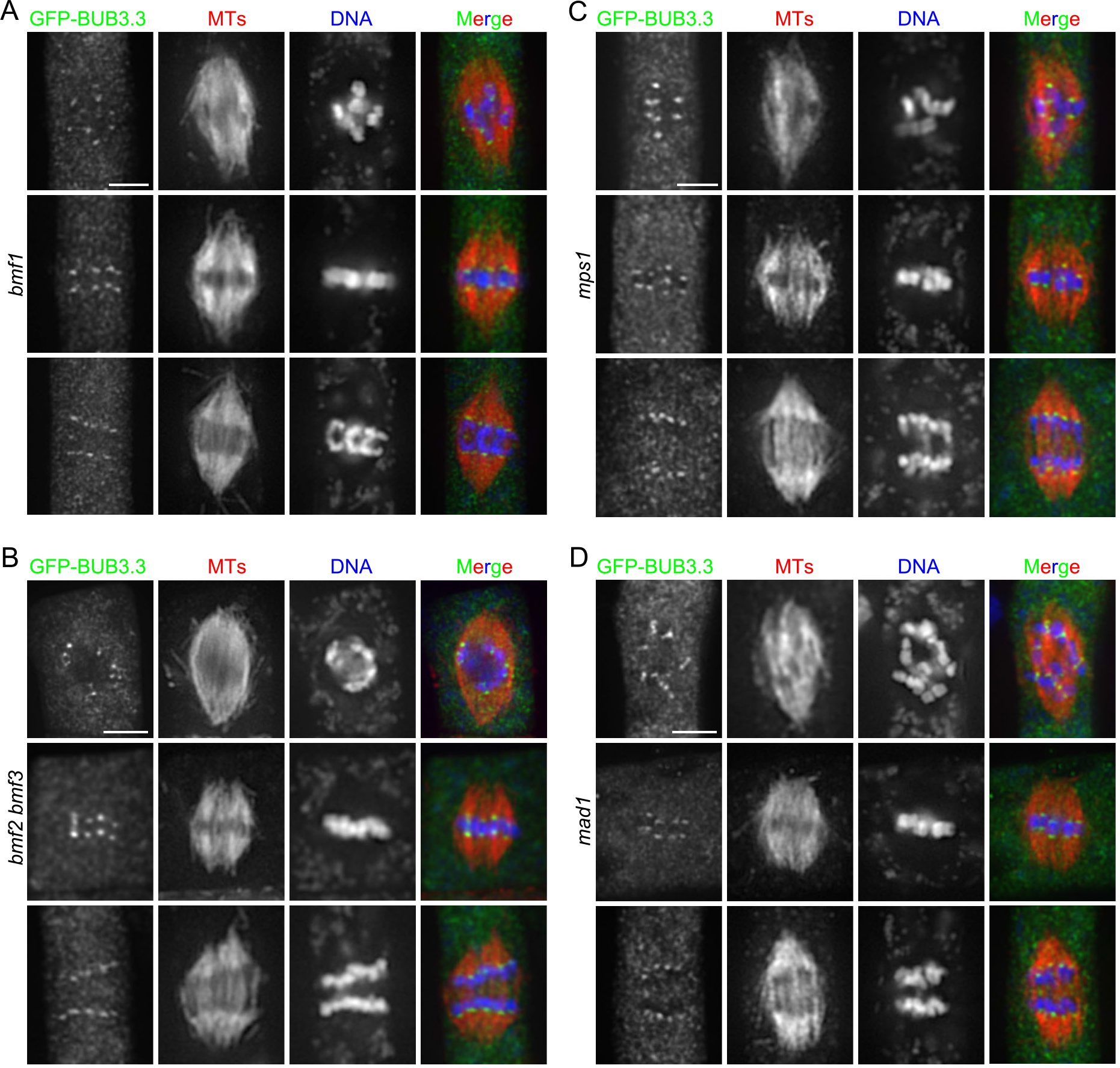
The kinetochore localization of BUB3.3 is independent to other SAC proteins. The GFP-BUB3.3 fusion protein was expressed and detected at kinetochores by immunostaining during mitosis in the *bmf1* (**A**), *bmf2 bmf3* (**B**), *mps1* (**C**) and *mad1* (**D**) mutant cells at prometaphase (top), metaphase (middle), and anaphase. Merged images have GFP-tagged proteins in green, microtubules in red, and DNA in blue. Scale bars = 5 μm.

### BUB3.3 recognizes two novel internal repeat motifs in BMF3

Because of the differential kinetochore localization patterns among different SAC components and their relationships, we then tested whether BUB3.3 interacted with one or more proteins described above. To do so, we employed yeast two-hybrid assays to examine potential interactions between BUB3.3 and BMF1, BMF2, BMF3, MAD1, and MAD2. We only detected the interaction between BUB3.3 and BMF3, but not others (Figure 6A). The two proteins also colocalized at kinetochores of the prometaphase chromosomes (Figure 6B), further supporting the notion that the two proteins were associated with each other *in vivo*.

**Figure 6.**
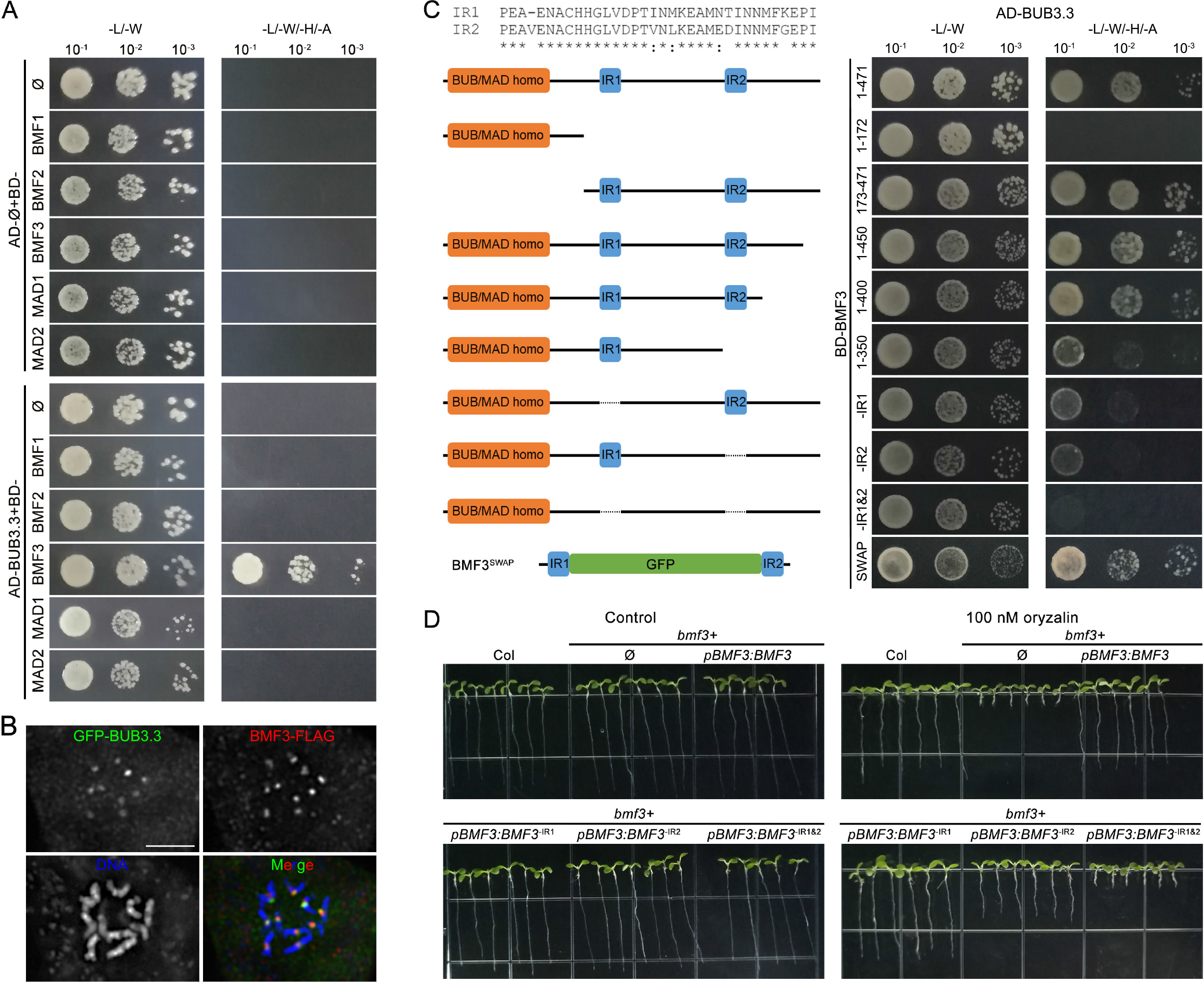
BUB3.3 recognizes two novel internal repeat (IR) motifs in BMF3. (**A)** Interaction between BUB3.3 interacts with BMF3 but not other Arabidopsis SAC components by the yeast two-hybrid (Y2H) assay. The empty vector is used as a negative control (Ø). The yeast cultures are spotted on vector-selective (-L/-W, left column) and interaction-selective (-L/-W/-H/-A, right column) media and photographed after incubation at 30°C for 2 days. (**B)** Colocalization of GFP-BUB3.3 and BMF3-FLAG in a mitotic cell. The merged image has BUB3.3 pseudocolored in green, BMF3 in red, and DNA in blue. (**C)** Schematic representations of full-length and truncated versions of BMF3 used for mapping the BUB3.3-binding domains by Y2H assay. The IR1 (amino acids 196-228) IR2 (amino acids 354-387) peptides are highly identical to each other. The BMF3^SWAP^ fusion protein has the region between IR1 and IR2 replaced by GFP. Note the deletion of either IR1 or IR2 significantly reduces the interaction strength and BMF3^SWAP^ interacts with BUB3.3 similarly as the full-length protein. (**D)** Growth phenotypes of 7-day-old plants associated with the expression of various BMF3 derivatives on media lacking or including 100 nM oryzalin. Scale bars = 5 μm

To learn how the BUB3.3-BMF3 interaction was established, we made truncations of BMF3 for the yeast two-hybrid assays. First, we separated BMF3 into two parts, the N-terminal BUB1/MAD3 characteristic TPR domain (BMF3^1-172^) and the remaining fragment with two internal repeats (IR1 and IR2) (BMF3^173-471^). It was found that the BMF3^173-471^ fragment was sufficient for the interaction with BUB3.3 (Figure 6C). We started to truncate the BMF3 protein from the C-terminus and found that deletions of peptides after the IR2 motif did not affect the interaction (Figure 6C). When IR2 was removed in BMF3^1-350^, however, the interaction was weakened albeit not abolished (Figure 6C). This suggested that IR2 likely played a role in the interaction. Therefore, we made a truncated BMF3 with the IR1 deleted and found that this BMF3^ΔIR1(196-288)^ fragment also exhibited a partially compromised interaction with BUB3.3 (Figure 6C). A similar result was recapitulated when the truncated BMF3^ΔIR2(354-387)^ fragment without the IR2 motif was used (Figure 6C). Simultaneous deletions of both IR1 and IR2 motifs completely abolished the interaction (Figure 6C). Therefore, the result suggested that these two internal repeats were required for the interaction. Then we asked the question whether the two IR repeats were sufficient for BMF3 interaction. To do so, we designed an artificial IR1-GFP-IR2 fusion protein and found that this BMF3^SWAP^ protein interacted with BMF3 in yeast cells (Figure 6C). We then examined the *in vivo* functionality of the aforementioned BMF3 truncations after having them expressed under their native promoter in the *bmf3* homozygous mutant background. The *bmf3* mutant grew indistinguishably from the wild-type control under unchallenged conditions but exhibited hypersensitivity to 100 nM oryzalin (Figure 6D), similar to what has been reported previously (4). We found that the BMF3^Δ196-288^ version without the IR1 motif was largely functional as the full-length protein and the BMF3^Δ354-387^ fragment without IR2 partially suppressed the bmf3 mutation (Figure 6D). The removal of both IR1 and IR2 abolished the functionality of BMF3 in this oryzalin challenge assay (Figure 6D). Therefore, we concluded that both IR1 and IR2 contributed to the interaction with BUB3.3 but IR2 probably was more critical than IR1 for the functionality of BMF3 when the SAC is turned on.

### The BUB3.3-BMF3 interaction activates the recruitment of CDC20 to the unattached kinetochores

Because BUB3.3 interacted with BMF3 but did not affect its kinetochore localization, we then examined whether other targets of BMF3 were affected by the interaction with BUB3.3. We first screened BMF3 binding proteins among MCC components via yeast two-hybrid assay. Indeed, BMF3 interacted with MAD1 (Figure 7A), as previously reported (4). We also found BMF3 interacted with CDC20.1 but not MAD2 by the yeast two-hybrid assay (Figure 7A). Because the localization of MAD1 was not affected by BUB3.3 as demonstrated above, we then examined whether CDC20.1 localization was affected by the loss of BUB3.3.

**Figure 7.**
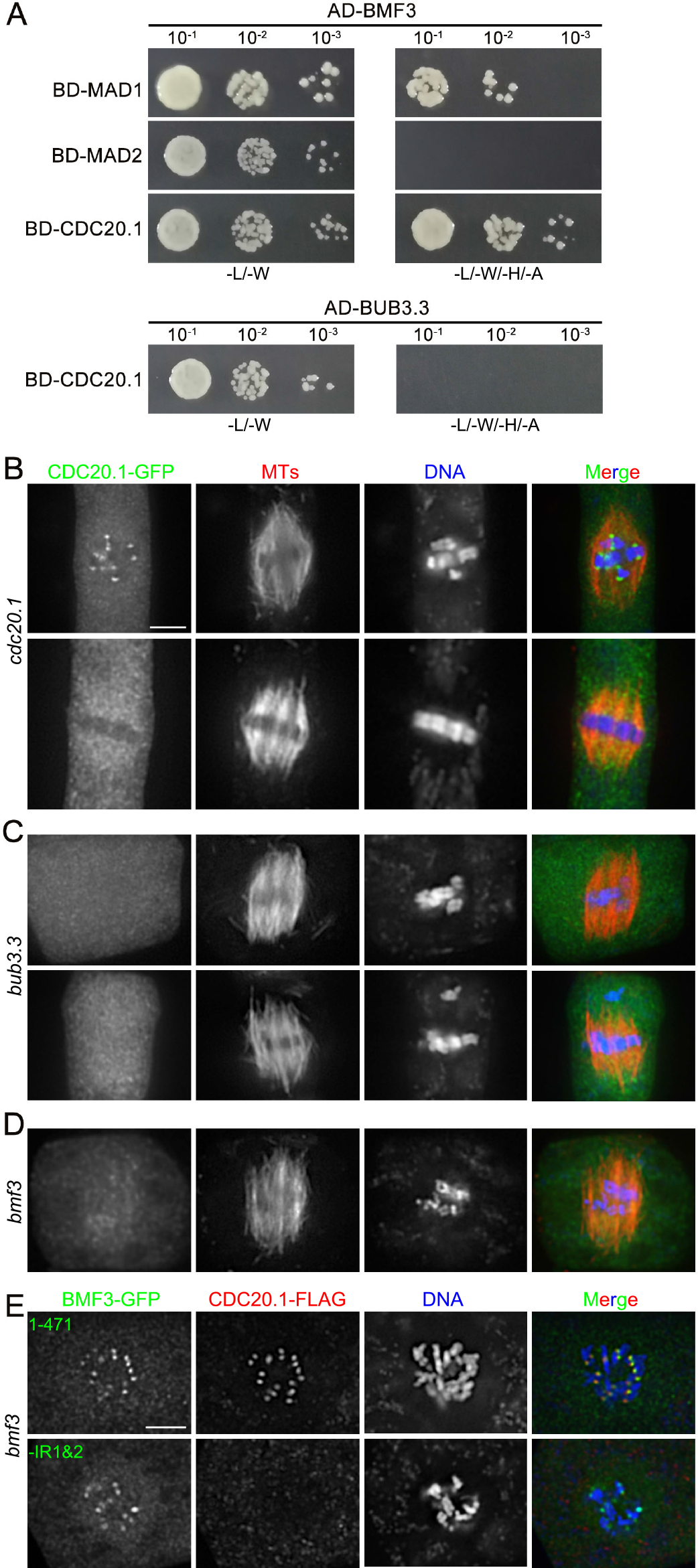
BUB3.3-BMF3 interaction is required for the recruitment of CDC20 to unattached kinetochores. (**A)** BMF3 interacts with MAD1 and CDC20.1 in a Y2H assay. But BUB3.3 does not interact with CDC20.1. (**B-D)** CDC20.1-GFP localization in different genetic backgrounds. When it is expressed in in the *cdc20.1* mutant (**B**), the fusion protein is detected at kinetochores of all chromosomes at prometaphase and becomes cytosolic upon chromosome congression at the metaphase plate. In the *bub3.3* (**C**) and *bmf3* (**D**) mutant cells, however, CDC20.1-GFP is no longer detected at kinetochores at any stages of mitosis. (**E)** When the CDC20.1-FLAG fusion protein is expressed in the bmf3 mutant cells co-expressing either the full-length BMF3-GFP (top rows) or truncated BMF3^ΔIR1&2^-GFP (bottom rows) fusion proteins, it colocalizes with BMF3-GFP at kinetochores but not BMF3^ΔIR1&2^-GFP which is still detected at kinetochores in prometaphase cells. In the merged images, the GFP fusion proteins are pseudocolored in green, microtubules by anti-tubulin (B-D) or CDC20.1-FLAG by anti-FLAG (E) in red, and DNA by DAPI in blue. Scale bars = 5 μm.

Because CDC20 is a crucial MCC component and the key target upon SAC activation and is structurally and functionally conserved in plants (1, 8), we then examined the CDC20.1 protein in mitotic cells after a CDC20.1-GFP fusion protein was expressed under the control of the *CDC20.1* promoter in the null *cdc20.1* mutant. CDC20.1-GFP was detected at kinetochores at prometaphase prior to chromosome congression to the metaphase plate (Figure 7B). The signal disappeared from there after chromosomes were amphitellicaly attached and aligned (Figure 7B). To determine whether such cell cycle-dependent localization was dependent on BUB3.3 and/or BMF3, the fusion protein was expressed in the *bub3.3* and *bmf3* mutants, respectively, after transformation. In the *bub3.3* mutant, the CDC20.1-GFP signal was no longer detected at kinetochores following nuclear envelope breakdown or those of misaligned chromosomes (Figure 7C). Similarly, CDC20.1-GFP was not at kinetochores either in the *bmf3* mutant (Figure 7D). Therefore, these results revealed that the kinetochore localization of CDC20 was a response to SAC activation and was dependent on both BUB3.3 and BMF3 in *A*. *thaliana*.

To further verify the significance of the internal repeats, we had the truncated BMF3^Δ(IR1+IR2)^ protein expressed in the *bmf3* mutant. Compared to the cells expressing the full-length BMF3-GFP which exhibited kinetochore association and restored the localization of CDC20.1 in a fusion with the FLAG tag, the mutant cells expressing the BMF3^Δ(IR1+IR2)^-GFP fusion protein no longer had CDC20.1-FLAG detected at kinetochores although the mutant BMF3 protein remained there (Figure 7E). These results collectively led to the conclusion that BUB3.3 did not govern the localization of BMF3 but was essential for activating BMF3 for the recruitment of CDC20 through the two novel internal repeats of BMF3.

## DISCUSSION

Our results showed a novel mode of action of the noncanonical BUB3 family protein BUB3.3 for its function in the activation but not the localization of BMF3 in order to recruit CDC20 to kinetochores of unattached chromosomes in *A*. *thaliana* as a reference system for higher plants. Our findings brought new insights into how plant cells respond to SAC activation by discovering a mechanism that employed the evolutionarily conserved BUB3 family protein to trigger SAC-dependent mitotic arrest without having to dissociate from the kinetochores to assemble equivalent of the MCC found in vertebrates.

### Novel BUB3.3-interaction domains in the plant-specific BMF3 protein

Because BUB3 family proteins are highly conserved and made of mostly six WD40 repeats, their interacting proteins often are said to contain the GLEBS domain which is responsible for the interaction and commonly found in BUB1/BUBR1 proteins in animal cells (6). However, the BMF proteins, which are distantly related to the animal BUB1 family proteins with isolated functional domain(s), lack this interaction motif, it has been unclear how BUB3.3 might be coupled with BMF family proteins to prevent anaphase onset upon the activation of the SAC (2, 4). In mammals, BUB3 also interacts with the BuGZ protein bearing the GLEBS domain and the kinetochore scaffold protein KNL1 by recognizing the phosphorylated MELT motif (9-11). However, the plant BMF proteins did not share significant sequence homology with these two mammalian proteins. Here we provided direct evidence showing that BUB3.3 interacted with two internal repeats found in the BMF3 protein. We did not find any vertebrate proteins that share significant sequence homology with these internal repeats. Therefore, the two repeats represent a new mode of interaction with the evolutionarily conserved BUB3 family proteins.

Among three BUB3 isoforms in *A*. *thaliana*, BUB3.1 and BUB3.2 are nearly identical and are more closely related to the animal or fungal BUB3 protein than BUB3.3 (12). Instead of functioning in SAC regulation, BUB3.1 and BUB3.2 directly interact with the microtubule-associated protein MAP65-3, localize to the phragmoplast midzone, and play redundant roles in regulating MAP65-3 function in phragmoplast microtubule reorganization during cytokinesis (4, 5, 13). The BUB3.3 shown here, however, exhibited exclusive localization at the centromeres and played an authentic role in SAC regulation, despite being more divergent from the fungal or animal BUB3 proteins. Collectively, these findings support the notion that the classical interaction modules of BUB3-GLEBS/MELT perhaps do not apply to flowering plants. Instead, plants established an alternative interaction module of BUB3.3-BMF3 internal repeats after BUB3 isoforms had their functions diversified.

### BUB3.3 exhibits SAC activation independent kinetochore localization

In human cells, BUB3 localization at kinetochores shows a drastic reduction from prometaphase to metaphase, suggesting a great deal of BUB3 delocalization from there after SAC activation (14). The kinetochore localization of BUB3.3, however, did not show obvious fluctuations throughout mitosis. In fact, different SAC-implicated proteins have different dynamic patterns during mitosis in *A*. *thaliana* and only BMF3 and MAD1 exhibit clear SAC activation-dependent localization patterns (4). Like BUB3.3, BMF1 and the key SAC kinase MPS1 also are associated with kinetochores throughout mitosis. Unlike the MAD2 homolog in maize which localizes to kinetochores of unaligned chromosomes, the Arabidopsis counterpart seems to be predominantly cytosolic (4, 15). These phenomena raised the questions of whether the functions of these proteins are converged towards SAC regulation and if so how these proteins are functionally linked in plant cells.

In animal cells, a key contribution of BUB3 is to recruit BUB1 and BUBR1/MAD3 to kinetochore (16). Such a dependency is not shown in plant cells as demonstrated by the BUB3.3-independent localization of MAD1 and BMF3 to the kinetochores of unattached or lagging chromosomes. Conversely, BUB3.3 also did not require BMF3 for its kinetochore localization. Such independence was further supported by the kinetochore localization of the truncated BMF3 lacking the internal repeats required for BUB3.3 interaction. These results raised a possibility that the function of BUB3.3 in SAC regulation may be separated from those of MAD1 and BMF3. This notion is supported by the phenotype of chromosome misalignment in the *bub3.3* mutant that was not observed in the *mad1* and *bmf3* mutants.

Our results also raised the question of how SAC proteins including BUB3.3 achieve kinetochore localization. In animal cells, the KNL1 protein presented the phosphorylated MELT motif to recruit BUB3 (6). Plants produce proteins that share homologies in two functional domains with animal KNL1 proteins but lack an obvious MELT motif and localize to kinetochore in mitotic and meiotic cells (3, 17, 18). Furthermore, the maize KNL1 homolog demonstrates direct interactions with the BMF1 and BMF2 homologs (3). Therefore, the classical KMN network is likely formed by KNL1 and the Mis12 and NDC80 complexes in plants (19). Although it has been shown that inactivation of the KNL1 gene in maize leads to defects in chromosome congression during mitosis (3), it is yet to be tested whether the plant KNL1 homologs are required for the kinetochore localization of either BMF1, BMF2, or BUB3 during mitosis.

### Inhibition of CDC20 upon SAC activation

The ultimate consequence of SAC activation is the inhibition of the APC/C activator CDC20 *via* the formation of the MCC complex as demonstrated in animal cells (16). In *A*. *thaliana*, the function of CDC20 in spindle assembly and chromosome segregation has been demonstrated in male meiotic cells (8). Here, we demonstrated direct interaction between BMF3 and CDC20.1 and BMF3 was required for CDC20.1 localization to kinetochores. Furthermore, BUB3.3 was required for CDC20.1 localization to kinetochores, although the kinetochore localization of BMF3 was not dependent on BUB3.3. These findings suggested to us that BUB3.3 perhaps functioned in the activation of BMF3 for the recruitment of CDC20.1 to kinetochores in order to catalyze the inhibition of APC/C and anaphase onset. On the one hand, this is different from animal cells in which BUB3 is required for BUB1 and BUBR1 localization to kinetochores (20). On the other hand, such an action may partly resemble the activation of BUBR1 by BUB3 in human cells in which BUB3 enhances the tethering of BUBR1 at kinetochores for CDC20 recruitment (21). It was unclear whether plant cells assemble the equivalent of the animal MCC complex and if so what such an MCC complex is made of (4). Among the three BMF proteins, BMF1 is the only one that bears a kinase domain but is likely dispensable for SAC because its mutant lacks the oryzalin hypersensitivity phenotype; and BMF2 is abundantly cytosolic but not most prominent at kinetochores (4). Therefore, MCC assembly in *A*. *thaliana* probably included BUB3.3, BMF3, and CDC20 but not BMF1 and BMF2 because of their different localization patterns. This notion was also supported by the data that BUB3.3 did not interact with BMF1 or BMF2 in our experiments.

However, the oryzalin hypersensitivity phenotype of the *bmf2* mutant and the predominant cytosolic localization of the BMF2 protein suggested to us that it might contribute to SAC signaling in the cytosol. However, such cytosolic SAC signaling scheme must be BUB3.3 independent, unlike the scenario in animal cells, again because BUB3.3 only interacted with BMF3 but not BMF2. It is interesting that MAD1 interacts with both MAD2 and BMF3 in *A*. *thaliana* (4). However, unlike MAD1 which exhibits a typical SAC activation-dependent kinetochore localization, MAD2 shows predominant cytosolic localization in addition to kinetochores which is dependent on MAD1. Because BMF3 is required for the kinetochore localization of MAD1, upon SAC activation, MCC might be assembled in the following sequential steps.

BMF3 was activated by BUB3.3 at kinetochores while recruited MAD1 which catalyzed the conformational change of MAD2. When CDC20 was recruited to unattached kinetochores by BUB3.3-activated BMF3, it then complexed with MAD2 in the closed conformation. The catalysis of MAD2-CDC20 complex formation perhaps dissociated from the unattached kinetochores and associated with BMF2 in the cytosol to form the MCC complex. Alternatively, MCC formation may be independent from the kinetochore localization. This hypothesis was supported by at least two lines of evidence generated in animal cells. For example, it was suggested that the interaction between BUB3 and CDC20, MAD2, and MAD3 does not require intact kinetochores (22). An independent work also led to the hypothesis that CDC20-MAD2 association may be established in a kinetochore-independent manner (23).

Furthermore, because of BUB3.3-independent localization of BMF3 and other SAC proteins, it would be interesting to learn whether the proposed kinetochore scaffolding factor KNL1, despite being divergent from the animal counterparts, plays a role in the localization of both BMF3 and BUB3.3. Interestingly, the maize KNL1 homolog interacts with BMF1 and BMF2 but not BMF3 in the plant, despite the fact that all three proteins possess the TPR domains (3). It is particularly intriguing that BMF3 localized to unattached kinetochores at prometaphase only while BUB3.3 had continuously appearance at all kinetochores. Such a difference suggested to us that the two proteins had different localization mechanisms.

### BUB3.3 and chromosome congression/alignment

We found that the loss of BUB3.3 led to frequent misalignment of chromosomes during mitosis and this chromosome congression phenotype was drastically enhanced when the cells experienced mild microtubule depolymerization challenges. As a result, micronuclei were born from the misaligned chromosomes because the *bub3.3* mutant cells entered anaphase prematurally. It is unclear whether this phenotype was directly linked to defects besides that of SAC signaling. In animal mitosis, congression of chromosome to the metaphase plate is dependent on various microtubule-associated factors, like the Kinesin-7 motor CENP-E cancer cells (24). In cancer cells, SAC activation following chromosome misalignment leads to micronucleus formation (24). There are 15 Kinesin-7 isoforms in *A*. *thaliana* (25), but to date it is unclear whether one or more of them function as CENP-E and whether they are functionally linked to SAC proteins like BUB3.3.

## MATERIALS AND METHODS

Plant materials, plant growth conditions, plasmid construction, yeast two-hybrid assays, microscopic observation, and imaging are described in SI Appendix, SI Materials and Methods.

## ACKNOWLEDGEMENT

This work was supported by the National Science Foundation (NSF) grant MCB-1920358 to YRJL and BL, and National Natural Science Foundation of China (32270354) and the Institutional Research Fund of Sichuan University (Grant No. 2020SCUNL212) to XD. BL is supported by from the U. S. Department of Agriculture (USDA)-the National Institute of Food and Agriculture (NIFA) under an Agricultural Experiment Station (AES) hatch project (CA-D-PLB-2536-H). We are very grateful to Prof. T. Nakagawa for sharing the pGWB vectors.

## SUPPORTING INFORMATION (SI)

### SI MATERIALS AND METHODS

### SUPPLEMENTARY FIGURE AND MOVIE LEGENDS

**SI Movie S1**. Live-cell imaging of GFP-BUB3.3 and mCherry-labelled microtubules in the *bub3.3* background. Images have intervals of 15 s.

**SI Movie S2**. Live-cell imaging of WT cells expressing NDC80-TagRFP following the treatment with 100 nM oryzalin. Images have intervals of 20 s.

**SI Movie S3**. Live-cell imaging of *bub3.3* cells expressing NDC80-TagRFP following the treatment with 100 nM oryzalin. Images have intervals of 20 s.

## Supporting Information (SI)

## SI MATERIALS AND METHODS

### Plant Materials and Growth Conditions

All T-DNA insertional lines of *A. thaliana* were obtained from the Arabidopsis Biological Research Center (ABRC) at Ohio State University. They were *bub3.3* (SALK_022904 at the AT1g69400 locus), *bmf1* (SALK_122554 at AT2g20635), *bmf2* (SAIL_303_E05 at AT2g33560), *bmf3* (SALK_032111 at AT5g05510), *mad1* (SALK_073889 at AT5g49880), *cdc20.1* (GABI_438H03 at AT4g33270) and *mps1* (GABI_663H07 at AT1g77720). All plants were grown in a controlled environment with 16-h-light and 8-h-dark cycle at 22°C. Seedlings for live-cell imaging and immunolocalization were produced on solid medium supplied with 1/2 Murashige Skoog (MS) and 0.8% phytagel.

For oryzalin sensitivity experiments, seeds were germinated on 1/2 MS solid medium supplied with 100 nM oryzalin in DMSO and equal volumes of DMSO served as the control. Ten-day-old seedlings grown on plates with or without oryzalin were photographed and root length was measured by ImageJ. For live-cell imaging, seedlings were germinated on 1/2 MS solid medium for 4 days, then transferred to plates containing either 100 nM oryzalin or DMSO for 12h.

### Plasmid Construction

Genomic fragments of each gene, which contain the promoter and coding sequences, were amplified and cloned into pDONR221, followed by LR recombination reactions with pGWB650 to yield a C-terminus GFP fusion, or pGWB10 to get a FLAG fusion. To create GFP-BUB3.3 construct, the Entry vector containing genomic BUB3.3 was linearized by inverse PCR, then a EGFP fragment was inserted in front of the start codon via Gibson Assembly method (New England Biolabs). The resulting entry clone were recombined with pGWB1 by LR clonase. The entry vectors containing CDS of MAD1, MAD2, BMF2 or BMF3, Pro_MAD1_:GFP:MAD1 and Pro_BMF3_:BMF3:GFP were described as previously. Primer pairs for plasmid construction are listed in SI Table S1.

### Yeast Two-Hybrid Assay

For yeast two-hybrid assay, all of the cDNAs tested were amplified and cloned into pDONR221. The subcloned cDNAs were recombined into the pGADT7-GW (AD) or pGBKT7-GW (BD) through LR recombination reactions. The resulting constructs were transformed into the yeast strain AH109 and were spotted on SD plates without Leu and Trp (-L/-W; control media) or without Leu, Trp, His and Ade (-L/-W/-H/-A; selection media) and photographed after incubation at 30°C for 2 days.

### Immunolocalization and Fluorescence Microscopy

Root meristematic cells from 5-day-old seedlings were prepared for immunofluorescence staining as described previously (1). Primary antibodies used in this study included GFP recombinant rabbit monoclonal antibody (ThermoFisher, Catalog G10362), DM1A mouse anti-α-tubulin monoclonal antibody (Abcam, Catalog ab7291) and 9A3 mouse anti-FLAG monoclonal antibody (Cell Signaling Technology, Catalog 8146). Secondary antibodies were Alexa fluor 488-conjugated goat anti-rabbit IgG and Alexa fluor 555-conjugated goat anti-mouse IgG (ThermoFisher, Catalog A32731 and A32727). Stained cells were observed under an Eclipse 600 microscope equipped with 60X Plan-Apo and 100X Plan-Apo objectives (Nikon). Images were acquired by an OptiMOS camera (Q Imaging) controlled by the µManager software package.

For live-cell observation, meristem cells of 5-day-old seedlings were observed under an LSM710 laser scanning confocal module (Carl Zeiss) with a 40X C-Plan (water) objective. Images were acquired using the ZEN software package (Carl Zeiss) and processed in ImageJ.

---

**SI Figure S1.**
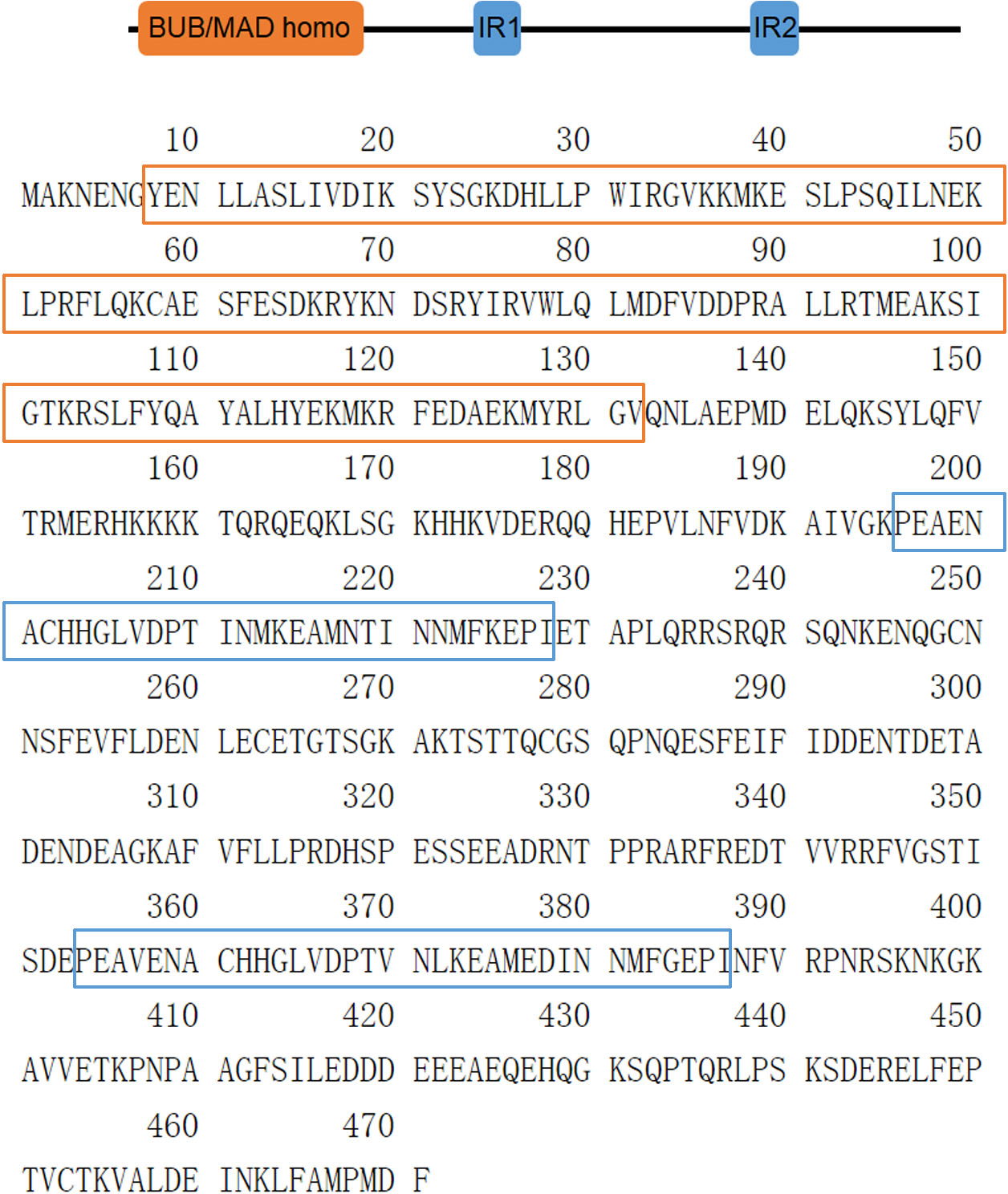
Schematic representations of the functional domains and corresponding sequences of the BMF3 protein in *Arabidopsis thaliana*. The BUB/MAD homology domain is highlighted by orange boxes and internal repeats (IR) are highlighted by blue boxes.

**SI Table S1.**
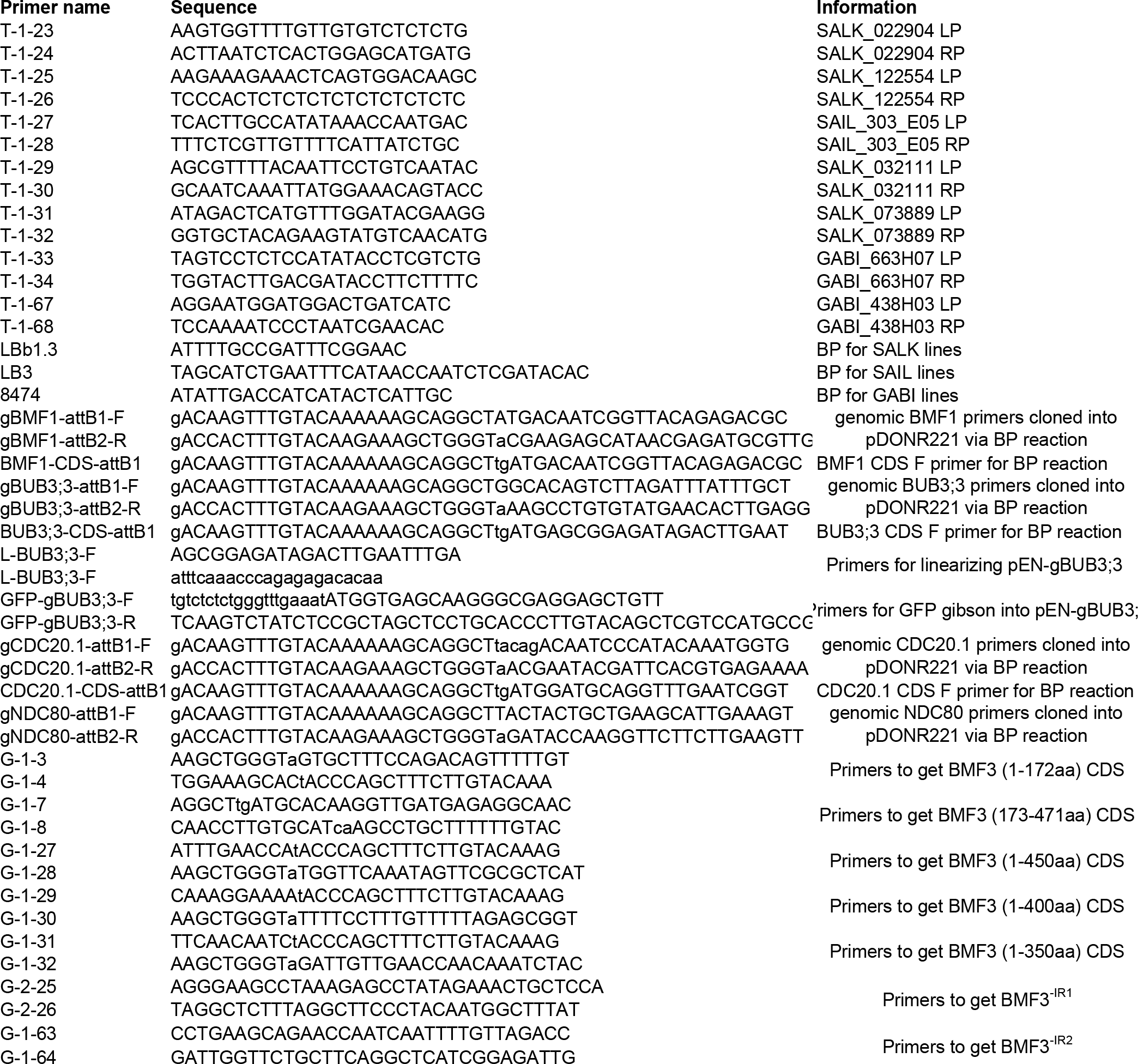
Primers used in this study.

## REFERENCES

1. Lara-Gonzalez P, Pines J, & Desai A (2021) Spindle assembly checkpoint activation and silencing at kinetochores. Seminars in Cell & Developmental Biology 117:86–98.

2. Komaki S & Schnittger A (2016) The spindle checkpoint in plants-a green variation over a conserved theme? Curr Opin Plant Biol 34:84–91.

3. Su H, et al. (2021) Knl1 participates in spindle assembly checkpoint signaling in maize. Proc Natl Acad Sci U S A 118(20).

4. Komaki S & Schnittger A (2017) The Spindle Assembly Checkpoint in Arabidopsis Is Rapidly Shut Off during Severe Stress. Dev Cell 43(2):172–185 e175.

5. Zhang H, et al. (2018) Role of the BUB3 protein in phragmoplast microtubule reorganization during cytokinesis. Nat Plants 4(7):485–494.

6. Kops G, Snel B, & Tromer EC (2020) Evolutionary Dynamics of the Spindle Assembly Checkpoint in Eukaryotes. Curr Biol 30(10):R589–R602.

7. Shin J, Jeong G, Park JY, Kim H, & Lee I (2018) MUN (MERISTEM UNSTRUCTURED), encoding a SPC24 homolog of NDC80 kinetochore complex, affects development through cell division in Arabidopsis thaliana. Plant Journal 93(6):977–991.

8. Niu BX, et al. (2015) Arabidopsis Cell Division Cycle 20.1 is required for normal meiotic spindle assembly and chromosome segregation. Plant Cell 27(12):3367–3382.

9. Jiang H, et al. (2014) A Microtubule-Associated Zinc Finger Protein, BuGZ, Regulates Mitotic Chromosome Alignment by Ensuring Bub3 Stability and Kinetochore Targeting. Developmental Cell 28(3):268–281.

10. Toledo CM, et al. (2014) BuGZ Is Required for Bub3 Stability, Bub1 Kinetochore Function, and Chromosome Alignment. Developmental Cell 28(3):282–294.

11. Vleugel M, et al. (2015) Sequential Multisite Phospho-Regulation of KNL1-BUB3 Interfaces at Mitotic Kinetochores. Molecular Cell 57(5):824–835.

12. Lermontova I, Fuchs J, & Schubert I (2008) The *Arabidopsis* checkpoint protein Bub3.1 is essential for gametophyte development. Front Biosci 13:5202–5211.

13. Caillaud MC, et al. (2009) Spindle assembly checkpoint protein dynamics reveal conserved and unsuspected roles in plant cell division. PLoS One 4((8)):e6757.

14. Shirnekhi HK, Herman JA, Paddison PJ, & DeLuca JG (2020) BuGZ facilitates loading of spindle assembly checkpoint proteins to kinetochores in early mitosis. J Biol Chem 295(43):14666–14677.

15. Yu HG, Muszynski MG, & Kelly Dawe R (1999) The maize homologue of the cell cycle checkpoint protein MAD2 reveals kinetochore substructure and contrasting mitotic and meiotic localization patterns. J Cell Biol 145(3):425–435.

16. Barisic M, Rajendraprasad G, & Steblyanko Y (2021) Spindle assembly checkpoint activation and silencing at kinetochores. Seminars in Cell & Developmental Biology 117:99–117.

17. Kozgunova E, Nishina M, & Goshima G (2019) Kinetochore protein depletion underlies cytokinesis failure and somatic polyploidization in the moss Physcomitrella patens. eLife 8.

18. van Hooff JJE, Tromer E, van Wijk LM, Snel B, & Kops GJPL (2017) Evolutionary dynamics of the kinetochore network in eukaryotes as revealed by comparative genomics. EMBO reports 18(9):1559–1571.

19. Liu B & Lee YRL (2022) Spindle assembly and mitosis in plants. Annu Rev Plant Biol 73:227–254.

20. Taylor SS, Ha E, & McKeon F (1998) The human homologue of Bub3 is required for kinetochore localization of Bub1 and a Mad3/Bub1-related protein kinase. Journal of Cell Biology 142(1):1–11.

21. Han JS, Vitre B, Fachinetti D, & Cleveland DW (2014) Bimodal activation of BubR1 by Bub3 sustains mitotic checkpoint signaling. Proc Natl Acad Sci U S A 111(40):E4185–4193.

22. Fraschini R, et al. (2001) Bub3 interaction with Mad2, Mad3 and Cdc20 is mediated by WD40 repeats and does not require intact kinetochores. EMBO J 20(23):6648–6659.

23. Li J, Dang N, Wood DJ, & Huang JY (2017) The kinetochore-dependent and -independent formation of the CDC20-MAD2 complex and its functions in HeLa cells. Scientific reports 7:41072.

24. Gomes AM, et al. (2022) Micronuclei from misaligned chromosomes that satisfy the spindle assembly checkpoint in cancer cells. Current Biology 32(19):4240-+.

25. Richardson DN, Simmons MP, & Reddy AS (2006) Comprehensive comparative analysis of kinesins in photosynthetic eukaryotes. BMC Genomics 7(1):18.

## REFERENCE

1. Lee YRJ & Liu B (2000) Identification of a phragmoplast-associated kinesin-related protein in higher plants. Curr Biol 10(13):797–800.

